# A new approach to crop model calibration: phenotyping plus post-processing

**DOI:** 10.1101/605220

**Authors:** Pierre Casadebaig, Philippe Debaeke, Daniel Wallach

## Abstract

Crop models contain a number of genotype-dependent parameters, which need to be estimated for each genotype. This is a major difficulty in crop modeling. We propose a hybrid method for adapting a crop model to new genotypes. The genotype-dependent parameters of the model could be obtained by phenotyping (or gene-based modeling). Then field data for example from variety trials could be used to provide a simple empirical correction to the model, of the form a+b times an environmental variable. This approach combines the advantages of phenotyping, namely that the genotype-specific parameters have a clear meaning and are comparable between genotypes, and the advantages of fitting the model to field data, namely that the corrected model is adapted to a specific target population. It has the advantage of being very simple to apply, and furthermore gives useful information as to which environmental variables are not fully accounted for in the initial model. In this study, this empirical correction is applied to the SUNFLO crop model for sunflower, using field data from a multi-environment trial network. The empirical correction reduced mean squared error (MSE) on the average by 54% for prediction of yield and by 26% for prediction of oil content, compared to the initial model. Most of the improvement came from eliminating bias, with some further improvement from the environmental term in the regression.

## Introduction

Crop model predictions depend on the model equations, the values of the input variables and the values of the parameters. Of course all three components are critical, but arguably it is the parameter values that present the greatest difficulty for model users. We focus here on parameter estimation for crop models.

Crop models often have a large number of parameters, perhaps more than 100. The STICS model for instance includes 132 parameters (Ruget et al., 2002). It is common to divide them into categories of generality. Some parameters are assumed to apply very generally and so are not meant to be altered for different applications. For example, in the SUNFLO model (Casadebaig et al., 2011; Lecoeur et al., 2011), there are parameters representing the effect of temperature and soil moisture on N mineralization rate, which are treated as fixed for all uses. Another set of parameters is specific to a particular species, which applies to generic models like DSSAT (Jones et al., 2003), APSIM (Holzworth et al., 2014) or STICS (Brisson et al., 2003) which can simulate for multiple species. For example, DSSAT has species senescence parameters related to vegetative stage, freeze damage, nitrogen remobilization, drought and canopy self-shading which are included in a species file. Finally, crop models have genotype-dependent parameters. For example, DSSAT uses six genotype-dependent parameters for maize, including photoperiod sensitivity and potential grain number. The SUNFLO model has 10 genotype-dependent parameters (two for degree days to key development stages, four for shoot architecture, two for response to water deficit and two for biomass allocation). For most model users, the general and species-specific parameters can be subsumed into *model structure*, since they are not altered from the values provided by the model developers. However, the problem is rather different for genotype-dependent parameters. There are very many varieties, which are often region-specific and furthermore new varieties are regularly developed and released. Obtaining genotype-dependent parameters is a never-ending problem.

There are three quite different approaches to estimation of genotype-dependent parameters that have been proposed. The most common approach (*estimation*) is to calibrate the model using field data for the variety in question, by searching the parameter space for the genotype-dependent parameters that give a good fit to the field data. An example is the study in Li et al. (2015) in which DSSAT maize and wheat are calibrated for new varieties (i.e. varieties not previously studied with DSSAT) in China. One disadvantage of this approach is that parameters are only available after the variety is disseminated, which may lead to a delay of several years between the development of the variety and the corresponding calibration of a crop model. In addition, in the absence of a standardized protocol, it may be difficult to compare varieties using a crop model, because differences in parameter values may also be due to differences in the type of data used for calibration. At the opposite extreme is gene-based modeling, where parameters are predicted as a function of the allelic composition of the genotype (*prediction*). In this case, models can be parameterized for new genotypes even before the genotype has been cultivated in the field. Developing such models is a major topic in modeling (Messina et al., 2006, 2018; Cooper et al., 2016; Hammer et al., 2016; Wallach et al., 2018), but as yet this approach is not widespread. An intermediate approach in terms of timing is model calibration based on phenotyping input parameters (*measurement*). In this approach, the genotype-dependent parameters of a crop model are measured using a standardized protocol of field or controlled environment experimentation coupled with detailed phenotype measures (e.g. Debaeke et al., 2010; Casadebaig et al., 2016b). This approach allows the measurement of genotype-dependent parameters at an early stage of variety testing. The fact that a standard protocol is used reduces the uncertainty in comparisons between varieties. High-throughput phenotyping could make this approach more efficient and allows one to include additional genotype-dependent parameters (Furbank and Tester, 2011; Cooper et al., 2014; Tardieu et al., 2017; Gosseau et al., 2019).

There are fundamental differences between the *estimation* approach and the *measurement* approach. The first one considers the model outcome, i.e. it estimates parameters values that can best describe the data, while the *measurement* approach considers physiological processes, by measuring parameters values that may or may not improve the prediction accuracy of model outcomes. The *measurement* approach also assumes that the genotype-dependent parameters, or more generally all the model parameters, can be applied in all environments. There is no mechanism in this approach to adapt the model to a specific set of environmental conditions, i.e. to a target population of environments. On the other hand, the calibration done in the *estimation* approach adjusts the model not only for the new variety, but also *de facto* adjusts the parameters to the target population represented by multi-environment trials. The difference would not be important, if indeed the same parameters were applicable to all environments, without need of adjustment depending on target population. However, it seems clear that the model is needed to be adapted to different target populations. A general argument is made by Jørgensen and Fath (2011) regarding models in ecology. Wallach et al. (2011) shows explicitly, for crop models, that calibrated parameters compensate for errors in fixed parameters, and that the compensation depends on the target population. A first conclusion is that the genotype-dependent parameters found by calibration to field data are not the “true” parameters, i.e. the values that one would obtain by studying each individual process of the crop model individually, and in the limit of a very large amount of data. A second conclusion is that calibration is needed for each new target population. Because of the assumptions made in the model, in the parameters selected for parameter estimation, error in input data, environment and management bias in the training sets, and limited ability to directly estimate all model parameters, local calibration can increase prediction accuracy.

Overall then, crop model calibration based on parameter estimation has the advantage that it adapts the model to a specific target population, but the disadvantage that the resulting genotype-dependent parameters are difficult to interpret and to compare between studies. Direct measurement of genotype-dependent parameters has the advantage that the parameters have physiological meaning, but the disadvantage that the model is not adapted for a specific target population. In gene-based models, because allelic composition is indeed constant between environments, the adaptation to a specific target population is moved to model structure.

In climate science, statistical post-processing techniques have emerged to provide a quantitative reinterpretation of raw numerical weather prediction models outputs, based on meteorological observations (Mendoza et al., 2015). Among these approaches, model output statistics (MOS) methods (Glahn and Lowry, 1972; Carter et al., 1989) have been typically used to develop forecast equations using dynamical model outputs and local observations as predictors.

The purpose of this study is to propose an hybrid approach which combines measurement of genotype-dependent parameters with calibration using field data. In this approach, similarly to MOS methods for weather prediction models, the calibration is done not by modifying the model parameters, but by adding a simple empirical adjustment to the model of the form a + b times an environmental variable. The goal is to combine the advantages of each parameterization approach; the measurement step produces genotype-dependent parameters of a crop model that have physiological meaning and the empirical adjustment produces a model that is adapted to a specific target population. We note that the field data needed for the systematic application of an empirical adjustment is often available, in the form of variety trials by breeders and/or post-registration multi environment trials (METs) by extension services.

We apply our proposed approach to the SUNFLO crop model. The model was specifically developed to use parameters that can be estimated by a standard phenotyping protocol, and has been parameterized in this way for a large number of sunflower varieties (Debaeke et al., 2010; Casadebaig et al., 2016b). The empirical correction is based on the large multi-environment trials conducted each year by breeders, examination offices and agricultural extension services in France to compare genotypes and assess their value for cultivation and use.

## Material and methods

### Field data

To train the linear model used in the calibration process, we relied on the variety testing network targeting released genotypes in France (Mestries and Jouffret, 2002) and conducted by the French technical institute in charge of references for oilseed crops and grain legumes (namely *Terres Inovia*). While this testing is conducted yearly, data is not in a readily usable format, and a major effort is needed to gather climate and soil data relevant to the tested locations. Here, we reused a dataset fully described in Casadebaig et al. (2016a), and summarized its main characteristics below. The target population represented by this dataset is sunflower cropping area in France with current recommended management.

Trials were conducted in 52 locations during one year (2009). Locations concentrating the most trials represented about 75 % of the cultivated sunflower area in France, i.e. Poitou-Charentes (16 trials), Centre (9), Midi-Pyrénées (8) and Pays de Loire (7) regions. The network was split in three geographical zones, matching the operational areas of the technical institute and referred to as South (South and South-West France), West (Center and West France), and East (North and East France). At each location, one to four variety trials were conducted (differing by maturity group or fatty acid composition – oleic vs linoleic), for a total of 82 trials performed over the network. Only the locations that could be reasonably described (nearby weather station, sufficient information on soil depth, reliable information on crop management) were kept in the dataset (82 out of 99 trials in the MET). Furthermore, only genotypes with data from at least 10 trials were retained (28 genotypes). In each trial, grain yield (*t.ha*^−1^, 0% humidity) and oil concentration (%, 0% humidity) were measured. Overall, the data represent 628 average plots (averaged over 3-4 replicates per genotypes × location modality)

### SUNFLO crop model description

SUNFLO is a process-based simulation model for sunflower that was developed to simulate grain yield and oil concentration as a function of time, environment (soil and climate), management practices and genetic diversity (Casadebaig et al., 2011; Lecoeur et al., 2011). Predictions with the model are restricted to attainable yield (Van Ittersum and Rabbinge, 1997): only the main limiting abiotic factors (temperature, light, water and nitrogen) are included in the algorithm.

The model simulates the main soil and plant functions: root growth, soil water and nitrogen dynamics, plant transpiration and nitrogen uptake, leaf expansion and senescence, and biomass accumulation. The proportion of biomass allocated to seeds (harvest index) and seed oil concentration are predicted with linear models based on variables simulated by the process-based model. Globally, the SUNFLO crop model has about 50 equations and 64 parameters split in 33 species parameters, 10 genotype-dependent, and 21 environment-related. A report that summarizes the equations and parameters used in the model is available as supplementary information in Picheny et al. (2017).

The values of the genotype-dependent parameters were obtained by measuring the value of the corresponding 10 phenotypic traits in dedicated field platforms and controlled conditions (table 1). Our aim was to measure potential trait values, so different environmental conditions were targeted depending on the set of traits: field non-limiting conditions (deep soil) for phenological and architectural traits, field pre-anthesis limiting conditions (shallow soil) for allocation traits, and a range of controlled water deficits (greenhouse) for response traits. For field experiments, sunflower hybrids were phenotyped in ten trials (two locations, five years: 2008-2012), using randomized complete block designs with three repetitions of 30 m^2^ plots (6-7 plant m^−2^), see Debaeke et al. (2010) and Casadebaig et al. (2016a) for trait measurement protocols. For controlled conditions, hybrids were phenotyped in 10 liters pots, during six greenhouse experiments to determine the response of leaf expansion and transpiration at the plant scale after stopping watering and letting the soil progressively dry (dry-down design). We used randomized complete block designs with two water treatments (control, stress) and six repetitions (7 pots m^−2^), see Casadebaig et al. (2008) for measurement protocol.

**Table 1:** Description and variation range for genotype-dependent parameters of the SUNFLO model. The maximum and minimum values reported represent the genotypic variability observed among phenotyped sunflower cultivars. Phenological traits were measured on the whole microplot (50% of the population required to reach the stage). Architectural traits were measured of 5 plants per microplot. Potential harvest index was estimated on 10 plants per plot, as the ratio of seed to shoot biomass, including senescent organs. Potential seed oil content was determined as the 9th decile of the distribution of oil concentration values measured in national networks for cultivar evaluation. Protocols for trait measurement are detailed in Casadebaig et al. (2016a).

| Function | Name | Description | Unit | Min | Max |
| --- | --- | --- | --- | --- | --- |
| Phenology | TDF1 | Temperature sum from emergence to the beginning of flowering | $Cd$ | 744.25 | 906.60 |
| Phenology | TDM3 | Temperature sum from emergence to seed physiological maturity | $Cd$ | 1460.80 | 2054.90 |
| Architecture | TLN | Potential number of leaves at flowering | *leaf* | 22.20 | 36.67 |
| Architecture | LLH | Potential rank of the plant largest leaf at flowering | *leaf* | 12.30 | 23.20 |
| Architecture | LLS | Potential area of the plant largest leaf at flowering | $cm^{-2}$ | 139.49 | 670.00 |
| Architecture | K | Light extinction coefficient during vegetative growth | - | 0.78 | 1.00 |
| Response | LE | Threshold for leaf expansion response to water stress | - | -15.57 | -2.15 |
| Response | TR | Threshold for stomatal conductance response to water stress | - | -14.21 | -5.51 |
| Allocation | HI | Potential harvest index | - | 0.25 | 0.51 |
| Allocation | OC | Potential seed oil content | % | 47.77 | 62.30 |

### Calibration process

Calibration was applied to the two output variables of the simulation model of major importance, namely crop grain yield and grain oil concentration.

First, the original model was run, and the model residuals were calculated. The model residuals are defined as *y_i_* – *f*(*X_i_*;*θ*) where *y_i_* is the measured value for environment *i, f*(*X_i_*; *θ*) is the corresponding simulated result which depends on the input variables *X_i_* and on the parameters *θ* obtained by phenotyping. The second step is to fit an empirical correction to the residuals, separately for each genotype.

In a first correction method (the *delta* method), the average of the residuals was calculated, i.e. the bias correction across environments, noted *â*. The corrected model is *f*(*X_i_*; *θ*) + *â*. In the second method (the *linear* method), the models *a* + *bZ_i_* were fit to the residuals, where *Z_i_* is one from a set of candidate input variables. The input variable which gave the smallest mean squared error (MSE), *Z_s_* was selected by comparing regression models. The model with linear correction is then 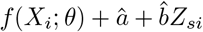 where *â* and 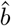 are the estimated regression coefficients.

A large number of input variables could be used for this empirical correction if we consider combinations of measured climatic variables by key crop phenological stages. Our proposal was to restrict this choice to a smaller number of candidates to limit selection bias. We therefore focused on variables readily available in climate datasets and representing the major abiotic factors known to affect sunflower growth and development. The four candidate input variables considered for the empirical correction are shown in table 2. They represent variables related to water (total precipitation or average water stress), radiation and temperature. In each case, a sum over the growing season, from sowing to maturity as simulated by SUNFLO, was used. For water deficit, the main abiotic stress in sunflower, we considered using a variable simulated by the model which was previously showed strong relation to observed yield (Mangin et al., 2017).

**Table 2.** Candidate variables for fitting SUNFLO residuals. Sums are over the growing season, from sowing to maturity as simulated by SUNFLO.

| label | factor | description |
| --- | --- | --- |
| Precipitation | water | Sum of precipitations |
| Radiation | light | Sum of photosynthetically active radiation |
| Temperature | temperature | Sum of days with mean temperature less than 20 °C |
| Water deficit | water | Sum of days with simulated water deficit |

### Evaluation of the calibration process

We used classical goodness-of-fit metrics between simulations (*x*) and observations (*y*). The first is mean-squared-error 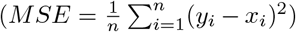, or root-mean-squared-error (RMSE) and the decomposition of MSE into a sum of three components (Kobayashi and Salam, 2000): squared bias 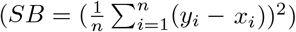, a term which measures the difference in spread between the observed and simulated values (Squared Difference between Standard Deviations, *SDSD* = *SD_x_* – *SD_y_*), and a term which depends on the correlation of measured and simulated values and so measures how well the simulated values track the variability in the observations (Lack of Correlation weighted by the Standard deviations, *LCS* = 2·*SD_x_* · *SD_y_* ·(1–*r*)).

Five-fold cross validation was used to evaluate the predictive accuracy of the corrected model. For each genotype, the data points were separated into five approximately equal parts (folds). Each fold in turn was designated the evaluation data, the remaining folds serving as training data. The linear correction, including the choice of the input variable for the empirical correction, was calculated using the training data and model prediction error was evaluated on the evaluation data. The procedure was repeated five times, so that each fold served once as evaluation data. In this way, each data point appeared exactly once in the evaluation data. The reported measures of predictive accuracy are averages over the five evaluation data sets.

### Software and data processing

Experimental and simulated data were processed with the R software version 3.5.1 (R Core Team, 2018) with R packages *dplyr* (data processing, Wickham et al., 2018), *rsample* (resampling, Kuhn et al., 2019), *ggplot2* (visualization, Wickham, 2016), and *knitr* (reporting, Xie, 2015). The source code for the SUNFLO simulation model is available on INRA software repository [https://forgemia.inra.fr/record/sunflo.git]. The INRA VLE-RECORD software environment (Quesnel et al., 2009; Bergez et al., 2013) was used as simulation platform.

## Results

### Environment-related input variables and relation to residuals

Figure 1 displays the variability in environment variables, over the 52 locations of the MET in 2009. Distributions of these variables tend to be bimodal, which can be explained by the spatial structure of the network, designed for testing late maturing cultivars in drought-prone locations (e.g. South-West of France), while early maturing cultivars were tested in colder and more humid environments. Because of this spatial variability most variables displayed a strong coefficient of variation (~ 35% for temperature and precipitation, ~ 60% for water deficit). Radiation showed lesser variability (~ 7%).

**Figure 1.**
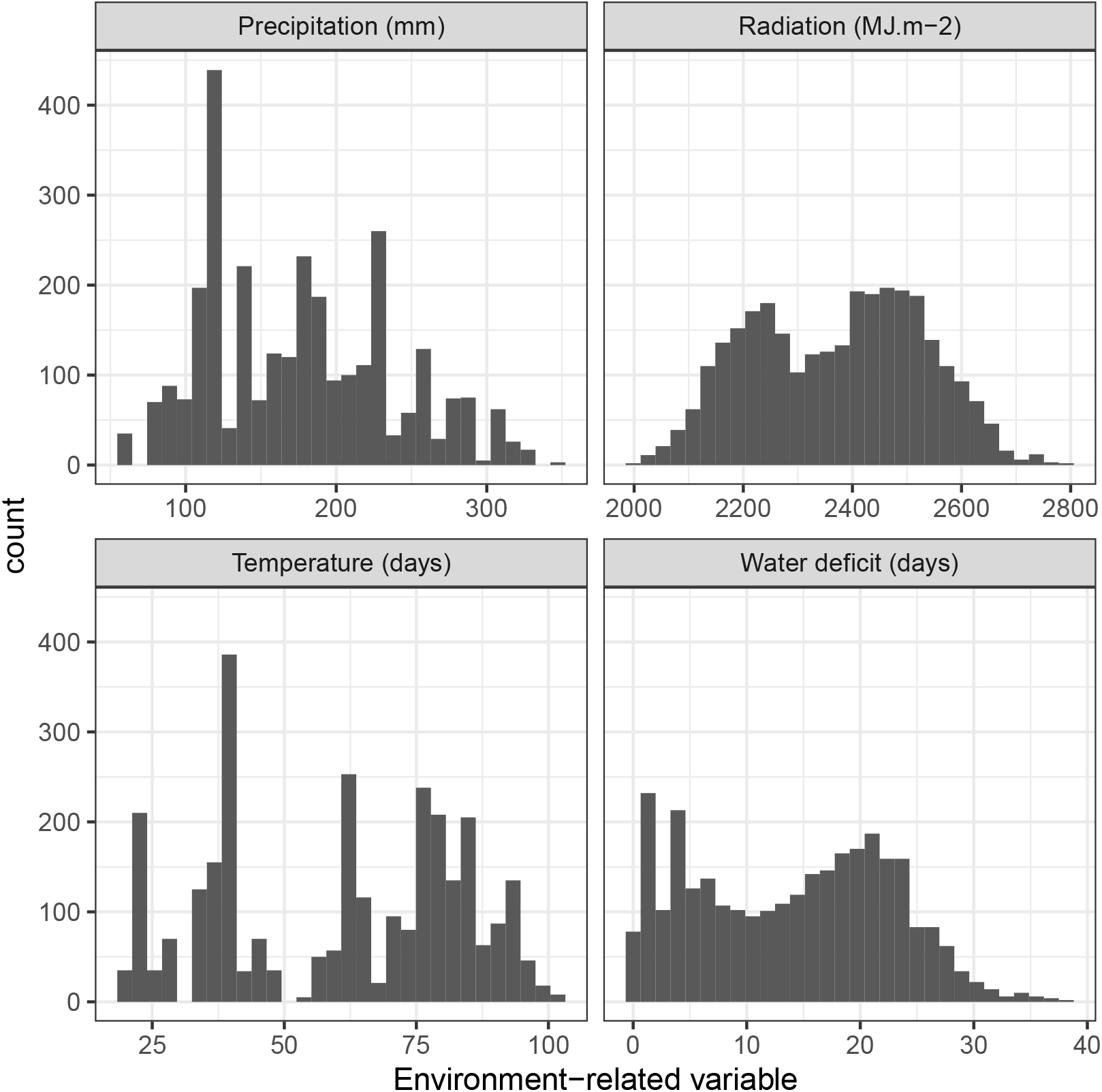
Distribution of selected environment-related variables in the network of trials. Each panel displays the histogram of selected environment-related variables: temperature (days with air temperature less than 20°C), water deficit (days with photosynthesis-limiting water deficit), radiation (sum of photosynthetically active radiation), and water (sum of precipitation). All sums are done over the growing season, from sowing to maturity as simulated by SUNFLO.

The regressions between the candidate variables and model residuals are presented in figure 2. Each panel only shows those genotypes for which the indicated candidate variable was chosen. Supplementary figure S1 illustrates these regressions in more detail for a subset of the dataset. Precipitation was the most-often chosen input to predict yield residuals (70 % of the population of genotypes) and temperature was most-often chosen for oil concentration residuals (55 % of the genotypes). For yield residuals related to precipitation, mean Pearson’s coefficient of correlation was 0.44 (ranging from −0.22 to −0.63). For oil concentration related to temperature, mean Pearson’s coefficient of correlation was –0.33 (ranging from −0.12 to −0.58). In two cases, a candidate input variable was not chosen for any genotype. Water deficit was not useful to explain grain yield residuals nor was radiation for oil concentration residuals.

**Figure 2.**
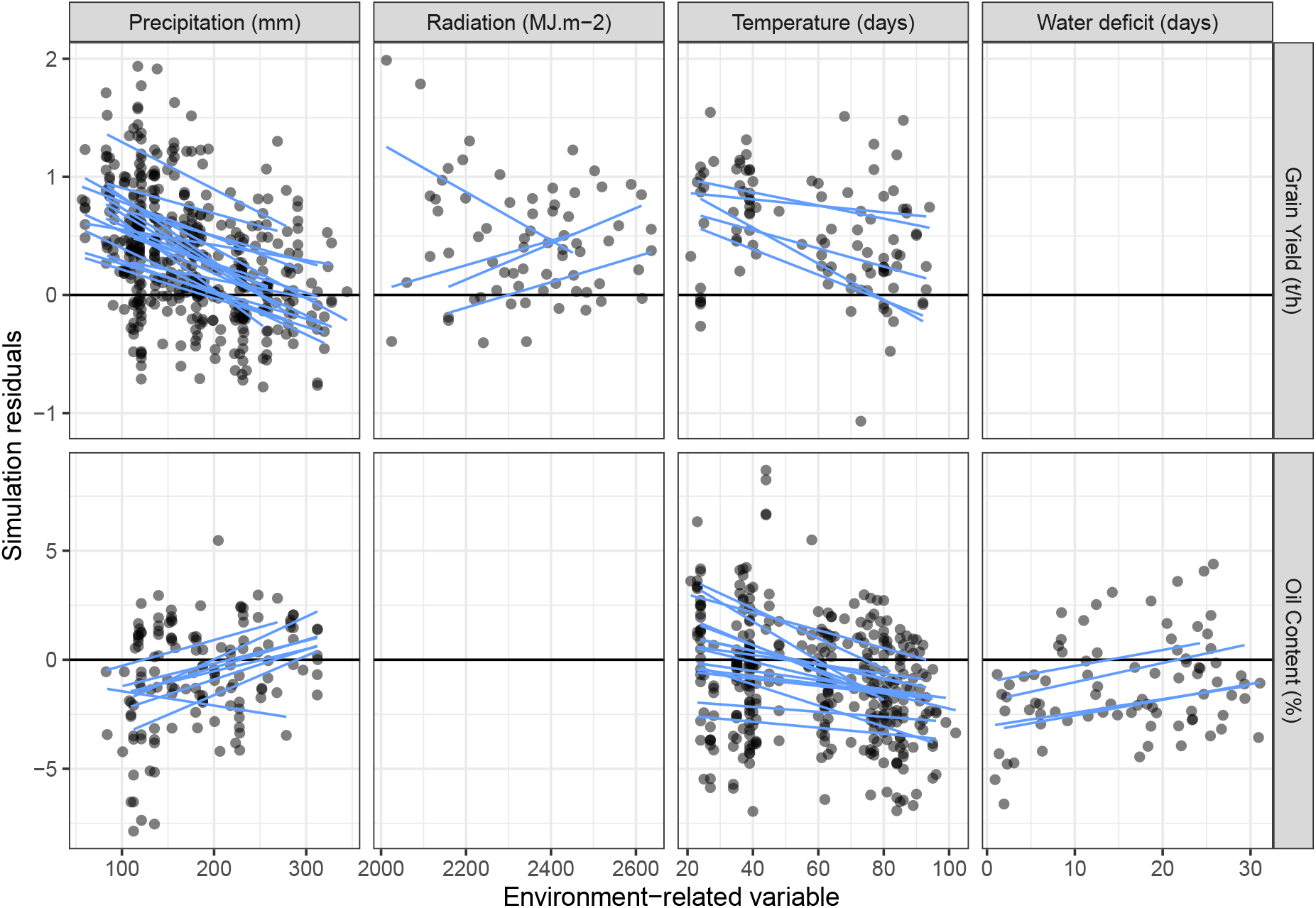
Linear relations between simulation model residuals and chosen environment-related variables. The panels display linear models fit to SUNFLO residuals (blue lines). Each line corresponds to one genotype. In each panel, data are only shown for those genotypes for which the indicated input variable gave the best fit. Output variables (Grain yield, oil concentration) are displayed in rows and environment-related variables are displayed in columns.

### Effect of empirical correction

The SUNFLO model with parameter values based on phenotyping, before any correction, has an average relative RMSE of 18% (0.63 t/ha) for yield and 5% (2.7%) for oil content (figure 3). The initial model accuracy on this dataset was slightly lesser than previously reported studies in Argentina or Spain (8-15%, Villalobos et al., 1996; Pereyra-Irujo and Aguirrezabal, 2007) or France (15%, Casadebaig et al., 2011). Even if the model correctly simulated phenotypic plasticity on this dataset and enabled applications for cultivar recommendation (Casadebaig et al., 2016a), it is clearly of interest to try to improve model predictions.

**Figure 3.**
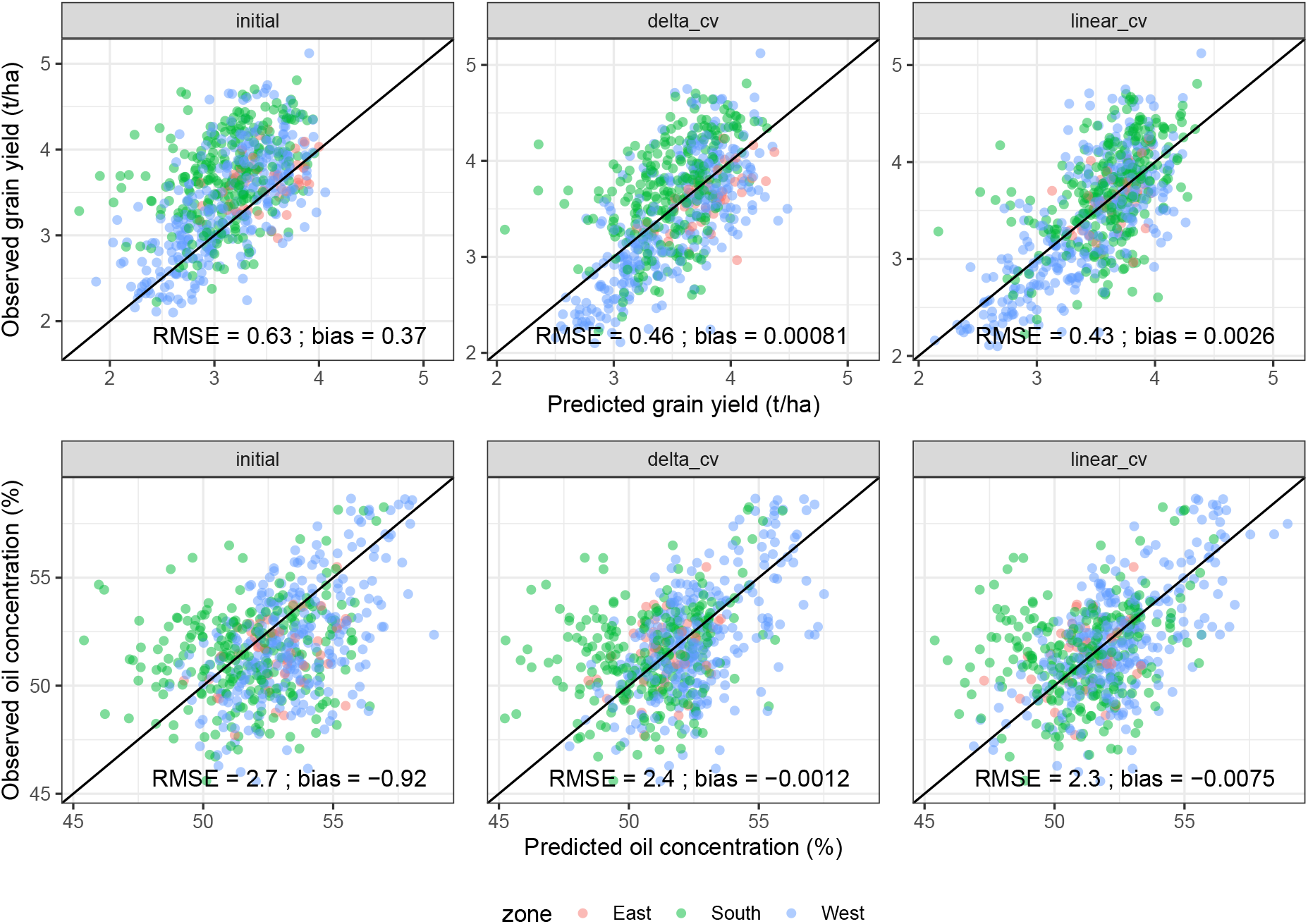
Observed vs predicted values after empirical correction. All genotypes are shown. Output variables are displayed in rows. The columns are the uncorrected model, the model corrected using just the average bias (delta method) and the model corrected using a linear function of the best input environmental variable (the linear method). The results are for cross validation.

We analyzed how the two empirical correction methods that we tested impacted the predicted values for grain yield and oil concentration (figure 3). Prediction quality was improved by the empirical correction process for all variables and methods, with the best improvement for the *linear* method. For grain yield correction, bias was essentially removed by the calibration methods and the linear regression further improved the predictions, particularly for the drought-prone cropping conditions (i.e. South zone in figure 3). This improvement was less important for oil concentration correction, with little additional improvement from the linear correction compared to the delta method. The model capacity to rank cultivars was also improved by empirical corrections: the Kendall’s *τ* improved from 0.36 to 0.46 for grain yield and from 0.25 to 0.34 for oil concentration.

We decomposed the mean-squared-error (MSE) into three error metrics (Kobayashi and Salam, 2000) to further analyse how each component was impacted by the calibration method (figure 4 and table 2).

**Figure 4.**
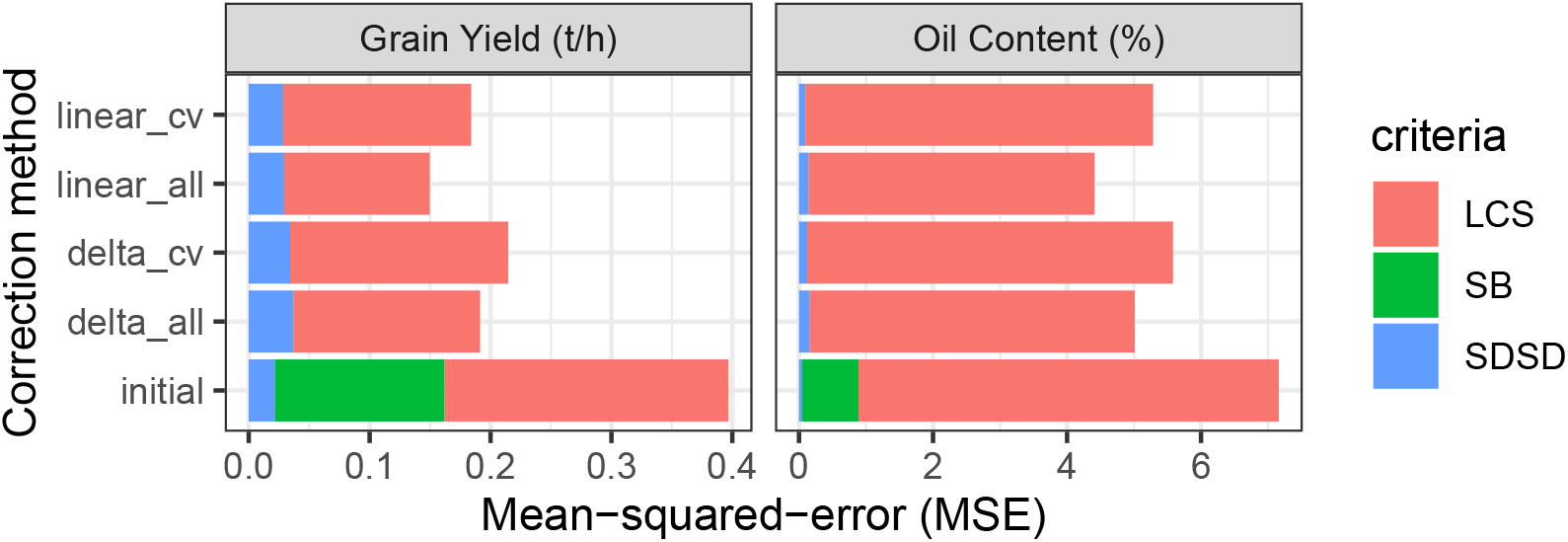
Components of mean-squared-error of prediction according to calibration methods. Results (bottom up) for the initial SUNFLO crop model (*initial*), after correction for just bias based on all the data (*delta_all*) and as estimated by cross validation (*delta_cv*), and after correction with the linear model based on all the data (*linear_all*) or estimated using cross validation (*linear_cv*).

About two thirds of MSE for yield for the initial model is due to the LCS term, which measures how well the simulated values track changes in the observations. About one third is from squared bias. A constant term in the linear correction will always eliminate bias exactly for the training data (*delta*). Most of that reduction still holds for prediction (as evidenced by the cross validation results). Adding an explanatory variable to the linear regression leads in addition to reduction in the LCS term, reducing it by about 34% for prediction (49 % in adjustment). The SDSD term, which measures the difference in the spread of the simulated values compared to the spread of the observations, is already very small for the initial model and remains small after correction.

The linear regression correction also eliminates the bias for oil concentration, both for the training data and for prediction. However, the squared bias is only 12% of MSE for the initial model, and so eliminating bias is relatively less effective for oil concentration than for yield. LCS is reduced by 17% by the linear regression correction, according to cross validation. Similar to yield, the SDSD term for oil concentration is very small initially and remains small.

**Table 3.**
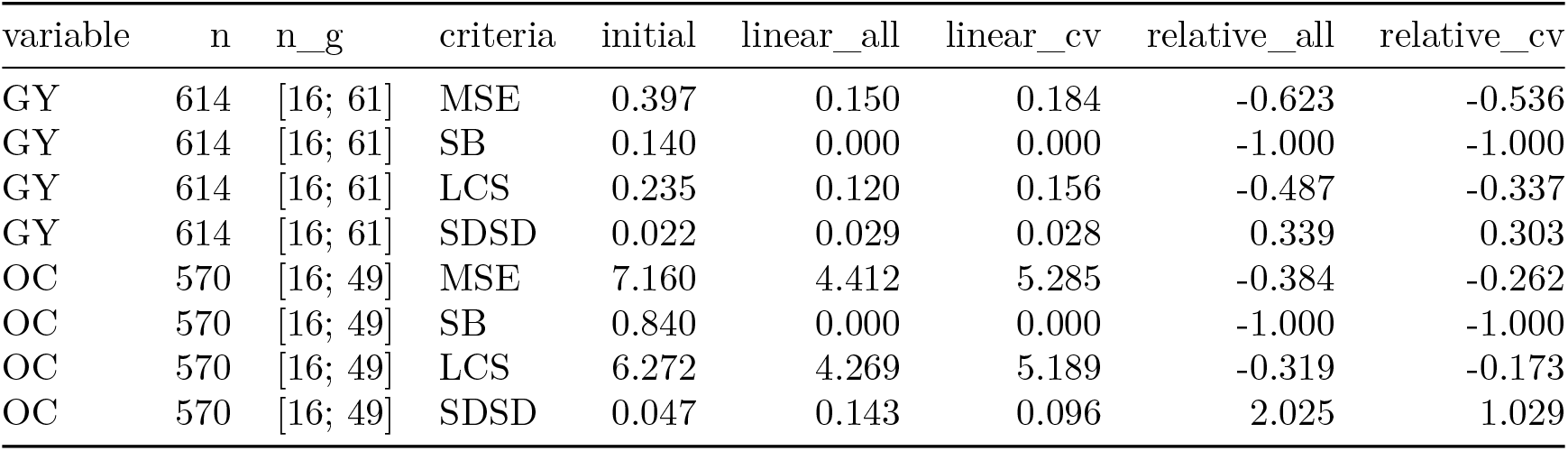
Components and relative mean-squared-error reduction after calibration. MSE and its components for grain yield (GY) or oil concentration (OC). The total number of observations (genotype × year combinations) is indicated in the *n* column, while the range of observations per genotype is indicated in the column *n_g*. The column *criteria* is MSE or one of the three components that add up to MSE. Results are for the initial SUNFLO model, after correction with the linear model based on all the data or estimated using cross validation. Relative differences (computed as (*corrected–initial*)/*initial*) between the corrected and initial models are shown in the *relative_all* and *relative_cv* columns.

| variable | n | n_g | criteria | initial | linear_all | linear_cv | relative_all | relative_cv |
| --- | --- | --- | --- | --- | --- | --- | --- | --- |
| GY | 614 | [16; 61] | MSE | 0.397 | 0.150 | 0.184 | -0.623 | -0.536 |
| GY | 614 | [16; 61] | SB | 0.140 | 0.000 | 0.000 | -1.000 | -1.000 |
| GY | 614 | [16; 61] | LCS | 0.235 | 0.120 | 0.156 | -0.487 | -0.337 |
| GY | 614 | [16; 61] | SDSD | 0.022 | 0.029 | 0.028 | 0.339 | 0.303 |
| OC | 570 | [16; 49] | MSE | 7.160 | 4.412 | 5.285 | -0.384 | -0.262 |
| OC | 570 | [16; 49] | SB | 0.840 | 0.000 | 0.000 | -1.000 | -1.000 |
| OC | 570 | [16; 49] | LCS | 6.272 | 4.269 | 5.189 | -0.319 | -0.173 |
| OC | 570 | [16; 49] | SDSD | 0.047 | 0.143 | 0.096 | 2.025 | 1.029 |

To investigate the sensitivity of our method to the number of data points, we analyzed how the correction efficiency (computed as the relative MSE reduction for each genotype) was impacted by the number of observations (i.e. environments) available for each genotype. The number of observations per genotype had a weak effect on the correction efficiency, with p-values for the hypothesis that there is no effect of 0.724 and 0.254 respectively for grain yield and oil concentration (supplementary figure S2).

## Discussion

The basic underlying hypothesis here is that there are no universally best parameter values for crop models. The best parameter values are somewhat different, depending on the target population. This is because models are not perfect; the parameter values compensate to some extent for inadequacies or errors in the model, and the required compensation depends on the target population. That is the reason that models need to be re-calibrated for different target populations. A statistical basis for this argument is given in Wallach et al. (2011).

The calibration strategy that is proposed here for the SUNFLO model is adapted to this situation. This is a strategy to improve the model for a particular target population. First, a standard protocol of phenotyping is used to estimate the genotype-dependent model parameters. It is expected, and found experimentally, that these parameters give fairly good model predictions for a range of environments. However, it is also expected that one can do better for a particular target environment by adapting the model to that specific environment. Thus the second stage of our strategy is to use a simple linear regression correction to adapt the model to a specific target population for which we have data. We note that both the phenotyping approach here and the second stage regression correction are for a specific genotype. Unlike gene-based models, this is not a methodology for extrapolating to new genotypes.

The strategy that is proposed is general; it can be used with the SUNFLO model whenever there is field data available, and it is specifically designed to be very easy to implement. However, the specific correction that is obtained here is not meant to be of universal validity. It is specifically meant to improve model predictions for the target population that provided the data here. A different correction would be required for a new set of data, from a different target environment. Since the correction is very simple, based on linear regression, it is quite easy to use cross validation to verify that the correction does improve predictions for the target population.

### Usefulness of an empirical correction

Our results show that the empirical correction we propose does provide a moderate improvement to model prediction, reducing MSE on the average over genotypes by 54% for prediction of yield and by 26% for prediction of oil content. This is done by making use of available variety testing data. To date, these data have been used simply to evaluate the SUNFLO model. The approach here shows how these data can be used for improving predictions, without fundamentally changing the initial model. The linear regression correction approach here is analogous to post processing for climate models, where simple statistical corrections are applied to GCMs, rather than tampering with the model parameters, to improve agreement with local weather data (Mendoza et al., 2015).

The effectiveness of the linear regression correction depends on the nature of the errors for the initial model. To the extent that bias is a major contribution to MSE of the initial model, the linear correction should be very effective since the constant term removes bias exactly for the training data. The results here show that adding a correction based on an explanatory variable makes the correction more effective by reducing the contribution of the LCS term, which is related to the correlation of the observed and simulated values, though it doesn’t completely eliminate the LCS contribution to MSE here. The results give no indication of whether the SDSD contribution to MSE, which measures the difference in variability among observed values compared to simulated values, will be effectively reduced, since this term plays only a negligible role here. While the prediction of absolute values and bias correction are important for applications in agronomy, the empirical corrections also improved the ranking of cultivars, which might be a more worthwhile model capacity for plant breeding.

In the absence of data to modify the model to account for causal effects on yield, the post processing method could be effective to expand the applicability of the model at reasonable cost. For example, on tomato in Florida, Messina et al. (2006) used a regression between model residuals and prices to calibrate their simulation model. The estimated parameters could thus be useful for a limited period and geographies.

The linear regression correction is clearly empirical. It is not meant as a method of improving the original model and original parameters. However, because the linear regression correction is based on relationships between simulation residuals and environment-variables it gives useful information that could help improve the initial model. The comparison of the original model and parameters with data, as always with model evaluation, gives information as to the level of model error. The procedure here goes a step further, and provides a systematic way of examining model residuals. Specifically, it is suggested to examine the residuals as a function of environmental variables likely to have an important impact on the system. These will not in general be the same variables as in the model, but rather more global variables. Here for example total rainfall is one of the variables considered for the regression correction, whereas the model depends on daily rainfall and daily evapotranspiration. The correction does not directly indicate how the model could be improved, but it does give indications as to possible problems.

In the present study, in most cases a correction related to precipitation improved the prediction of grain yield. Simulation error was better explained by a simple climate variable (rainfall) than a model state variable (water deficit). When focusing on the relation between observed yield (rather than residuals) and water deficit, we previously showed that model state variables were better predictors than climate variables (Mangin et al., 2017). For example, as seen in figure 1 the model tends to under predict grain yield (positive residuals) for low rainfall situations while the predictions are closer to observations (slightly negative residuals) for high rainfall environments. In this case, these observations suggest that while water deficit is accounted for (not correlated to simulation residuals but correlated to grain yield), some processes associated to rainfall are lacking in the SUNFLO model. Overestimation in high rainfall conditions could be related to disease development (e.g sclerotinia, phomopsis stem canker), usually occurring in wetter environments. Another possible explanation is that there are problems in estimation of soil water holding capacity, which has a greater impact in low rainfall environments than when more frequent rain will compensate for a poor soil water capacity estimation (indicated by lower residuals in high rainfall environments).

### Comparison with calibration of genotype-dependent parameters

A more common way of adjusting a crop model to data than the linear regression correction here is to start from some default parameter values and then to modify them to better fit the data. There is a large literature on crop model calibration done by modifying the parameters, and a large diversity of approaches (Seidel et al., 2018). Compared to linear regression, crop model calibration poses practical difficulties. Numerical problems arise because models are generally non continuous as functions of the parameters (Liu et al., 2018). Software difficulties arise in coupling a crop model to algorithms that search the parameter space. On the other hand, fitting a linear regression equation to model residuals is very simple, and execution time is not a problem.

More fundamentally, it has been argued that the calibrated parameter values of crop models are, at least to some extent, correcting for errors throughout the model, and that for different target populations, different values would be needed (Wallach et al., 2011). Therefore, at least to some extent, adjusting the parameters of the initial model to field data is not giving additional information about true phenotypic response, but also simply providing an empirical correction. Furthermore, there is some evidence that the major gain from crop model calibration is reduction in bias (Guillaume et al., 2011). The approach here on the other hand keeps the original, meaningful values of the parameters and is guaranteed to eliminate bias completely for the training data and robustly reducing it in test data. In addition, our linear regression correction is giving information on the causes of model error, whereas it is very difficult to interpret the results of modifying the initial parameter values.

### Possible improvements

The approach here uses a single environmental variable in the linear correction, chosen from among a small set of variables that are known to have a strong effect on yield and oil concentration. It is important to use knowledge of the crop to limit the number of possible input variables. With very many possible variables there is a high risk of selection bias, where random error rather than true correlation determines the choice of explanatory variable (Winship and Mare, 1992). When we tested this hypothesis on our dataset, goodness-of-fit was improved by allowing multiple explanatory variables, but prediction accuracy as evaluated by cross validation was decreased (figure S3). We assume that the tested linear models contained more parameters than can be justified by the data (overfitting). However, with more data, more explanatory variables could be envisioned.

A very promising improvement of the approach here would be to use a random parameter model. In this study, each genotype was treated separately. This has the disadvantage of limiting the available data for each correction to only the data for that genotype. In fact, it seems reasonable to suppose that the corrections for different genotypes are somewhat similar. That could be taken into account by using a model that treats the parameters of the linear regression as random variables, with some distribution, and the parameters for individual genotypes as being drawn from that distribution. This is known as “borrowing strength” (Steenbergen and Jones, 2002) and the effectiveness will depend on the number of data points available for each individual and how similar the regressions are between individuals. A limiting case of borrowing strength would be the case where no data are available for some genotypes. In that case, one could apply an empirical correction that uses the mean values of the parameters obtained for the available genotypes.

More generally, we argue that both process-based and data-based models are relevant tools to predict phenotypic plasticity. For exemple the function of the process-based model could be to simulate variables that reduce the dimension of climate data (multiple time-series) while accounting for dynamic plant-environment interactions (e.g. Messina et al., 2018, in the context of genomic prediction). In a second step, because field data at a large scale are available, data-based models might be relevant to predict complex traits (yield, quality, allocation) without having to explicitly model the related physiological processes, and by accounting for new limiting factors (e.g. the empirical relation between precipitation and yield probably accounts for biotic pressure).

## Conclusions

We propose a hybrid method for adapting a crop model to new genotypes. We suggest that the genotype-dependent parameters of the model could be obtained by phenotyping (or gene-based modeling). Then field data, especially variety trials, could be used to provide a simple empirical correction to the model. This approach combines the advantages of phenotyping, namely that the genotype-specific parameters have a clear meaning and are comparable between genotypes, and the advantages of fitting the model to field data, namely that the corrected model is adapted to a specific target population. It has the advantage of being very simple to apply, and furthermore gives useful information as to which environmental variables are not fully accounted for in the initial model. In this study, the empirical correction reduced MSE on the average by 54% for prediction of yield and by 26% for prediction of oil content. While the causal links are not fully understood and therefore included in the simulation model, this path could help to make crop models a part of broader and improved prediction models.

## Acknowledgements

Research grants were provided by the French National Research Agency (ANR SUNRISE ANR-11-BTBR-0005). The authors are grateful to the staff from *Terres Inovia* (Frédéric Bardy, Philippe Christante, André Estragnat, Pascal Fauvin, Emmanuelle Mestries, Jean-Pierre Palleau, Célia Pontet, Frédéric Salvi) that helped to constitute the phenotypic and climatic database.

## Supplementary material

**Figure S1.**
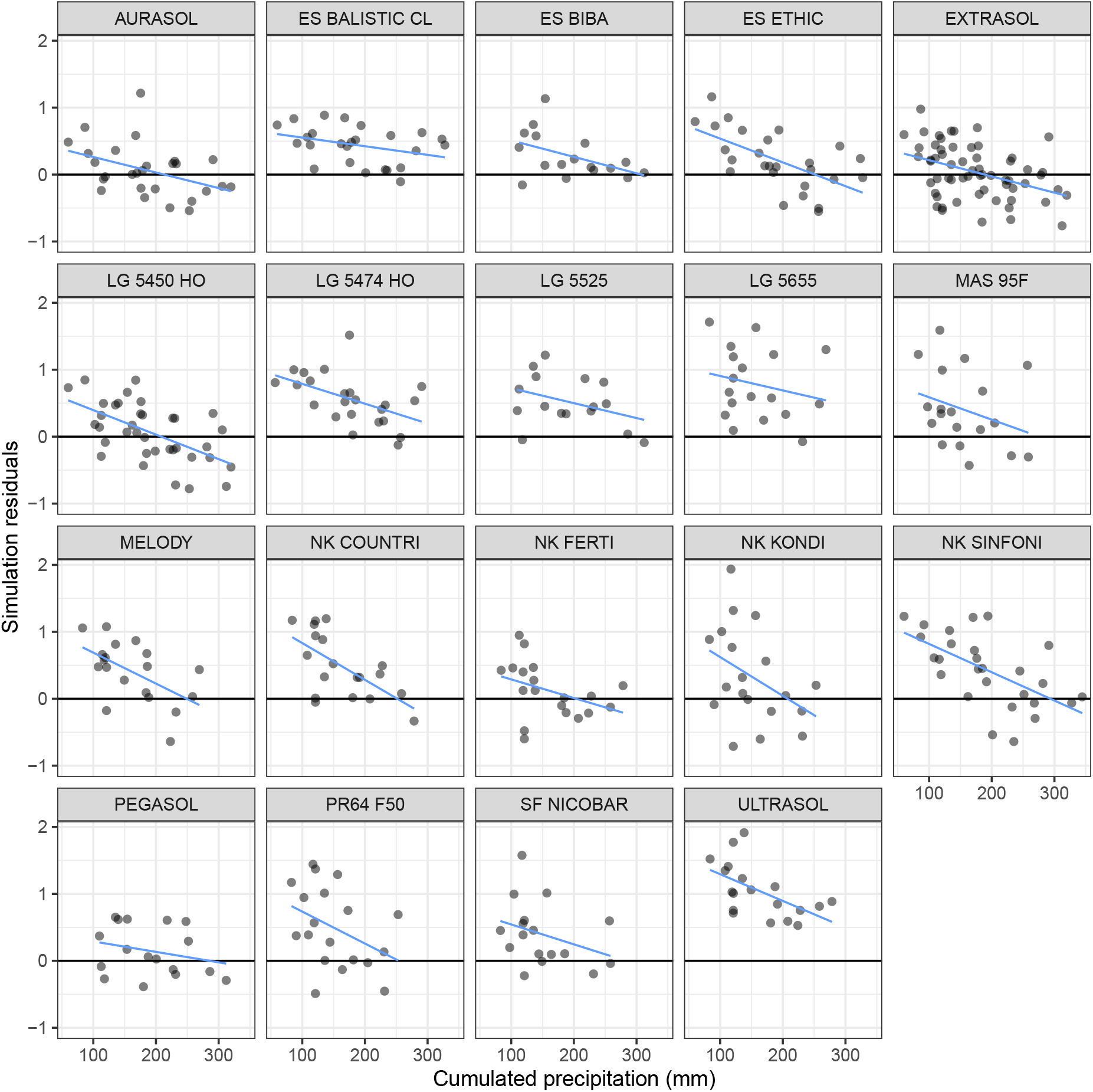
Linear regression of simulation model residuals versus selected environment-related variables. The panels display simulation model residuals as a linear function of accumulated precipitation, for each genotype with more than 10 observations in the trial network (blue lines). This figure shows the individual genotypes that are shown together in the upper left panel in figure 2.

**Figure S2.**
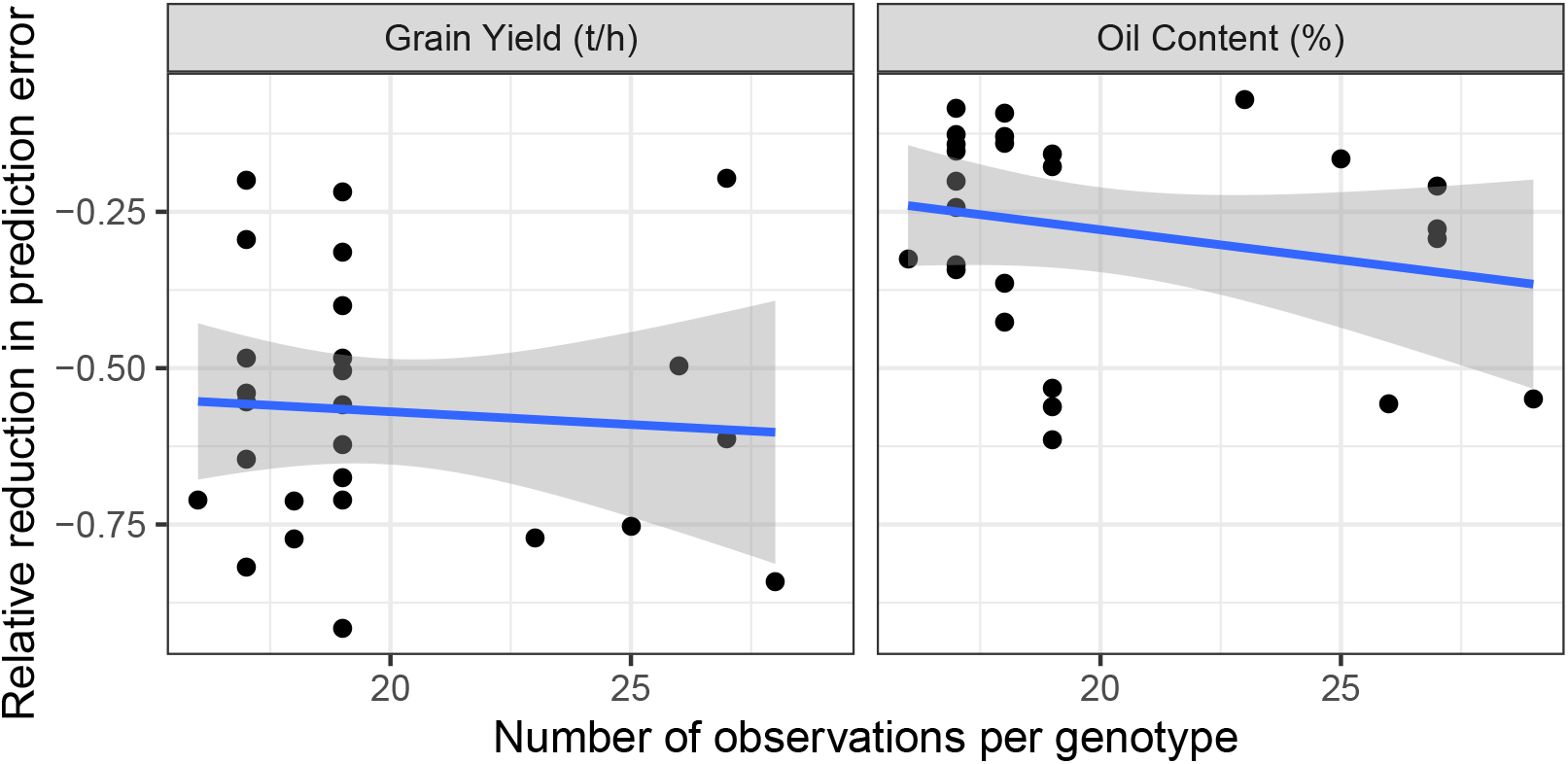
Efficiency of the calibration process as a function of the number of data points per genotype. The efficiency of the calibration process is calculated as the relative reduction in prediction error for each genotype ((*MSE_calibrated_ – MSE_initial_*)/*MSE_initial_*). Regression line is indicated in blue and uncertainty interval in grey.

**Figure S3.**
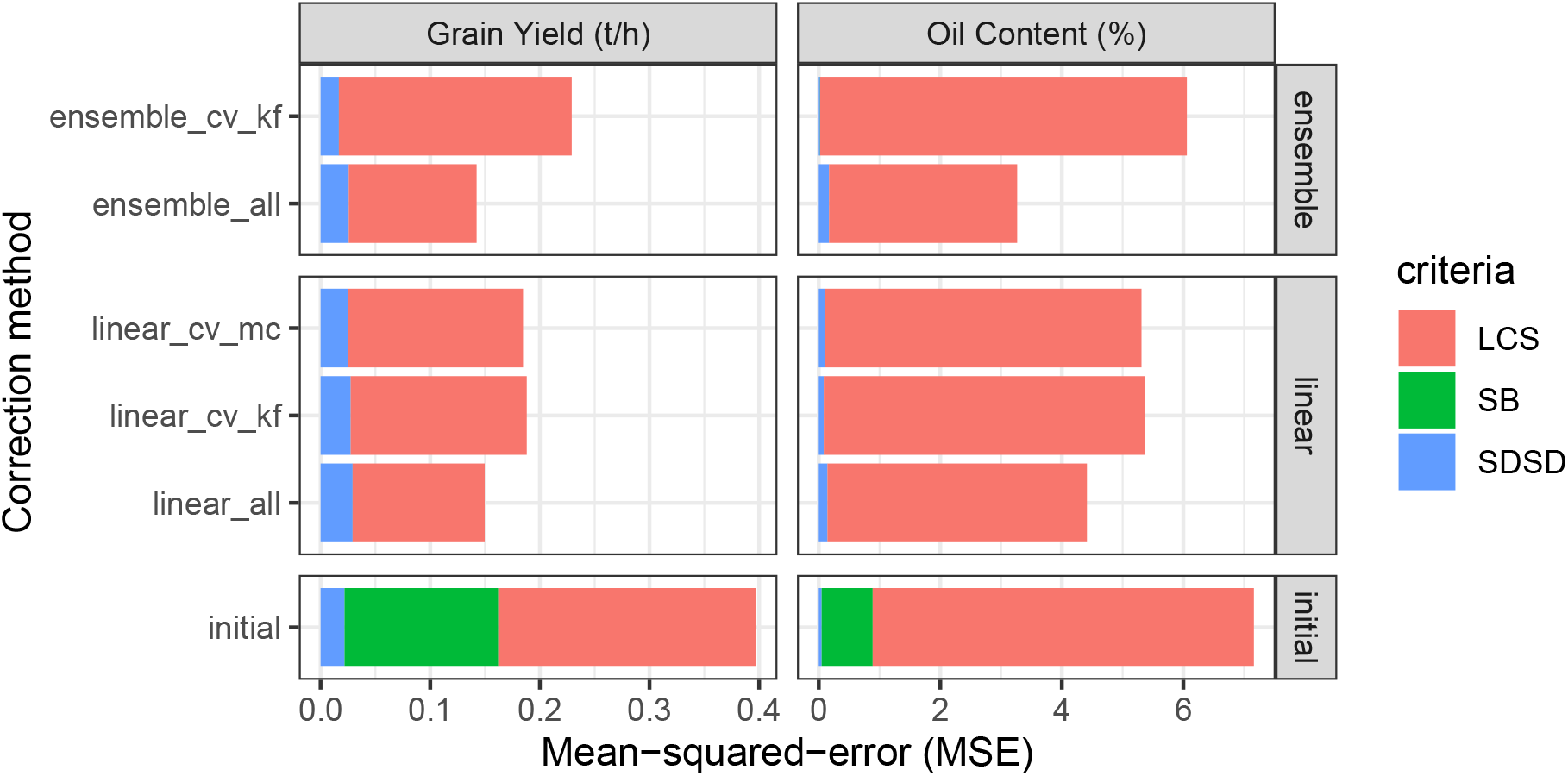
Components of mean-squared-error of prediction for calibration methods based on single or multiple variables. The decomposition of the prediction error is represented for three calibration options: the uncalibrated simulation model (*initial*), the calibration method with one explicative variable (*linear*), and with multiple explicative variables (*ensemble*). Three sampling strategies are also represented: using all the data to train the model (*all*), k-fold cross-validation (*cv_kf*), and monte-carlo cross-validation (*cv-mc*).

## References

Bergez, J., P. Chabrier, C. Gary, M. Jeuffroy, D. Makowski, et al. 2013. An open platform to build, evaluate and simulate integrated models of farming and agro-ecosystems. Environmental Modelling & Software 39: 39–49. doi: 10.1016/j.envsoft.2012.03.011.

Brisson, N., C. Gary, E. Justes, R. Roche, B. Mary, et al. 2003. An overview of the crop model STICS. European Journal of Agronomy 18(3-4): 309–332. doi: 10.1016/s1161-0301(02)00110-7.

Carter, G.M., J.P. Dallavalle, and H.R. Glahn. 1989. Statistical forecasts based on the national meteorological centers numerical weather prediction system. Weather and Forecasting 4(3): 401–412. doi: 10.1175/1520-0434(1989)004<0401:sfbotn>2.0.co;2.

Casadebaig, P., P. Debaeke, and J. Lecoeur. 2008. Thresholds for leaf expansion and transpiration response to soil water deficit in a range of sunflower genotypes. European Journal of Agronomy 28: 646–654. doi: 10.1016/j.eja.2008.02.001.

Casadebaig, P., L. Guilioni, J. Lecoeur, A. Christophe, L. Champolivier, et al. 2011. SUNFLO, a model to simulate genotype-specific performance of the sunflower crop in contrasting environments. Agricultural and Forest Meteorology 151: 163–178. doi: 10.1016/j.agrformet.2010.09.012.

Casadebaig, P., E. Mestries, and P. Debaeke. 2016a. A model-based approach to assist variety assessment in sunflower crop. European Journal of Agronomy 81: 92–105. doi: 10.1016/j.eja.2016.09.001.

Casadebaig, P., B. Zheng, S. Chapman, N. Huth, R. Faivre, et al. 2016b. Assessment of the potential impacts of wheat plant traits across environments by combining crop modeling and global sensitivity analysis (R. Wu, editor). PLOS ONE 11(1): e0146385. doi: 10.1371/journal.pone.0146385.

Cooper, M., C.D. Messina, D. Podlich, L.R. Totir, A. Baumgarten, et al. 2014. Predicting the future of plant breeding: Complementing empirical evaluation with genetic prediction. Crop and Pasture Science 65(4): 311–336. doi: 10.1071/cp14007.

Cooper, M., F. Technow, C. Messina, C. Gho, and L.R. Totir. 2016. Use of crop growth models with whole-genome prediction: Application to a maize multienvironment trial. Crop Science. doi: 10.2135/cropsci2015.08.0512.

Debaeke, P., P. Casadebaig, B. Haquin, E. Mestries, J.-P. Palleau, et al. 2010. Simulation de la réponse variétale du tournesol à l’environnement à l’aide du modèle sunflo. Oilseeds and fats, Crops and Lipids 17: 143–51. doi: 10.1684/ocl.2010.0308.

Furbank, R.T., and M. Tester. 2011. Phenomics technologies to relieve the phenotyping bottleneck.Trends in Plant Science 16(12): 635–644. doi: 10.1016/j.tplants.2011.09.005.

Glahn, H.R., and D.A. Lowry. 1972. The use of model output statistics (MOS) in objective weather forecasting. Journal of Applied Meteorology 11(8): 1203–1211. doi: 10.1175/1520-0450(1972)011<1203:tuomos>2.0.co;2.

Gosseau, F., N. Blanchet, D. Varès, P. Burger, D. Campergue, et al. 2019. Heliaphen, an outdoor high-throughput phenotyping platform for genetic studies and crop modeling. Frontiers in Plant Science 9. doi: 10.3389/fpls.2018.01908.

Guillaume, S., J.-E. Bergez, D. Wallach, and E. Justes. 2011. Methodological comparison of calibration procedures for durum wheat parameters in the STICS model. European Journal of Agronomy 35(3): 115–126. doi: 10.1016/j.eja.2011.05.003.

Hammer, G., C. Messina, E. van Oosterom, S. Chapman, V. Singh, et al. 2016. Molecular breeding for complex adaptive traits: How integrating crop ecophysiology and modelling can enhance efficiency. Crop systems biology. Springer. pp. 147–162

Holzworth, D.P., N.I. Huth, P.G. deVoil, E.J. Zurcher, N.I. Herrmann, et al. 2014. APSIM – Evolution towards a new generation of agricultural systems simulation. Environmental Modelling & Software 62(0): 327–350. doi: http://dx.doi.org/10.1016/j.envsoft.2014.07.009.

Jones, J.W., G. Hoogenboom, C. Porter, K. Boote, W. Batchelor, et al. 2003. The DSSAT cropping system model. European journal of agronomy 18(3): 235–265. doi: 10.1016/s1161-0301(02)00107-7.

Jørgensen, S.E., and B.D. Fath. 2011. Fundamentals of ecological modelling: Applications in environmental management and research. Elsevier.

Kobayashi, K., and M. Salam. 2000. Comparing simulated and measured values using mean squared deviation and its components. Agronomy Journal 92(2): 345–352. doi: 10.1007/s100870050043.

Kuhn, M., F. Chow, and H. Wickham. 2019. Rsample: General resampling infrastructure.

Lecoeur, J., R. Poiré-Lassus, A. Christophe, B. Pallas, P. Casadebaig, et al. 2011. Quantifying physiological determinants of genetic variation for yield potential in sunflower. SUNFLO: a model-based analysis. Functional Plant Biology 38(3): 246–259. doi: 10.1071/fp09189.

Liu, L., D. Wallach, J. Li, B. Liu, L. Zhang, et al. 2018. Uncertainty in wheat phenology simulation induced by cultivar parameterization under climate warming. European Journal of Agronomy 94: 46–53. doi: 10.1016/j.eja.2017.12.001.

Li, Z.T., J. Yang, C. Drury, and G. Hoogenboom. 2015. Evaluation of the DSSAT-CSM for simulating yield and soil organic c and n of a long-term maize and wheat rotation experiment in the loess plateau of northwestern china. Agricultural Systems 135: 90–104. doi: 10.1016/j.agsy.2014.12.006.

Mangin, B., P. Casadebaig, E. Cadic, N. Blanchet, M.-C. Boniface, et al. 2017. Genetic control of plasticity of oil yield for combined abiotic stresses using a joint approach of crop modeling and genome-wide association. Plant, Cell and Environment. doi: 10.1111/pce.12961.

Mendoza, P.A., B. Rajagopalan, M.P. Clark, K. Ikeda, and R.M. Rasmussen. 2015. Statistical postprocessing of high-resolution regional climate model output. Monthly Weather Review 143(5): 1533–1553. doi: 10.1175/mwr-d-14-00159.1.

Messina, C.D., D. Letson, and J.W. Jones. 2006. Tailoring management of tomato production to ENSO phase at different scales. Transactions of the ASABE 49(6): 1993–2003. doi: 10.13031/2013.22280.

Messina, C., F. Technow, T. Tang, R. Totir, C. Gho, et al. 2018. Leveraging biological insight and environmental variation to improve phenotypic prediction: Integrating crop growth models (cgm) with whole genome prediction (wgp). European Journal of Agronomy: –. doi: 10.1016/j.eja.2018.01.007.

Mestries, E., and P. Jouffret. 2002. Comment le CETIOM évalue les variétés. Oléoscope 66: 4–8.

Pereyra-Irujo, G.A., and L.A. Aguirrezabal. 2007. Sunflower yield and oil quality interactions and variability: Analysis through a simple simulation model. Agricultural and Forest Meteorology 143(3-4): 252–265. http://www.sciencedirect.com/science/article/B6V8W-4N094B0-1/2/0e62aeb380e0d4df21c1a643cd8a1429.

Picheny, V., P. Casadebaig, R. Trépos, R. Faivre, D. Da Silva, et al. 2017. Using numerical plant models and phenotypic correlation space to design achievable ideotypes. Plant, Cell & Environment. doi: 10.1111/pce.13001.

Quesnel, G., R. Duboz, and É. Ramat. 2009. The Virtual Laboratory Environment – An operational framework for multi-modelling, simulation and analysis of complex dynamical systems. Simulation Modelling Practice and Theory 17: 641–653. doi: 10.1016/j.simpat.2008.11.003.

R Core Team. 2018. R: A language and environment for statistical computing. R Foundation for Statistical Computing, Vienna, Austria.

Ruget, F., N. Brisson, R. Delécolle, and R. Faivre. 2002. Sensitivity analysis of a crop simulation model, STICS, in order to choose the main parameters to be estimated. Agronomie 22(2): 133–158. doi: 10.1051/agro:2002009.

Seidel, S.J., T. Palosuo, P. Thorburn, and D. Wallach. 2018. Towards improved calibration of crop models–where are we now and where should we go? European Journal of Agronomy 94: 25–35. doi: 10.1016/j.eja.2018.01.006.

Steenbergen, M.R., and B.S. Jones. 2002. Modeling multilevel data structures. American Journal of Political Science 46(1): 218. doi: 10.2307/3088424.

Tardieu, F., L. Cabrera-Bosquet, T. Pridmore, and M. Bennett. 2017. Plant phenomics, from sensors to knowledge. Current Biology 27(15): R770–R783. doi: 10.1016/j.cub.2017.05.055.

Van Ittersum, M., and R. Rabbinge. 1997. Concepts in production ecology for analysis and quantification of agricultural input-output combinations. Field Crops Research 52(3): 197–208. doi: 10.1016/s0378-4290(97)00037-3.

Villalobos, F., A. Hall, J. Ritchie, and F. Orgaz. 1996. OILCROP-SUN: A development, growth and yield model of the sunflower crop. Agronomy Journal 88: 403–415. doi: 10.2134/agronj1996.00021962008800030008x.

Wallach, D., S. Buis, P. Lecharpentier, J. Bourges, P. Clastre, et al. 2011. A package of parameter estimation methods and implementation for the STICS crop-soil model. Environmental Modelling & Software 26(4): 386–394. doi: 10.1016/j.envsoft.2010.09.004.

Wallach, D., C. Hwang, M.J. Correll, J.W. Jones, K. Boote, et al. 2018. A dynamic model with QTL covariables for predicting flowering time of common bean (phaseolus vulgaris) genotypes. European Journal of Agronomy 101: 200–209. doi: 10.1016/j.eja.2018.10.003.

Wickham, H. 2016. Ggplot2: Elegant graphics for data analysis. Springer-Verlag New York.

Wickham, H., R. François, L. Henry, and K. Müller. 2018. Dplyr: A grammar of data manipulation.

Winship, C., and R.D. Mare. 1992. Models for sample selection bias. Annual review of sociology 18(1): 327–350. doi: 10.1146/annurev.so.18.080192.001551.

Xie, Y. 2015. Dynamic documents with R and knitr. 2nd ed. Chapman; Hall/CRC, Boca Raton, Florida.

